# Non-Equilibrial Dynamics in Under-Saturated Communities

**DOI:** 10.1101/834838

**Authors:** Abdel Halloway, Kateřina Staňková, Joel S. Brown

## Abstract

The concept of the evolutionary stable strategy (ESS) has been fundamental to the development of evolutionary game theory. It represents an equilibrial evolutionary state in which no rare invader can grow in population size. With additional work, the ESS concept has been formalized and united with other stability concepts such as convergent stability, neighborhood invasion stability, and mutual invisibility. Other work on evolutionary models, however, shows the possibility of unstable and/or non-equilibrial dynamics such as limit cycles and evolutionary suicide. Such “pathologies” remain outside of a well-defined context, especially the currently defined stability concepts of evolutionary games. Ripa et al. (2009) offer a possible reconciliation between work on non-equilibrial dynamics and the ESS concept. They noticed that the systems they analyzed show non-equilibrial dynamics when under-saturated and “far” from the ESS and that getting “closer” to the ESS through the addition of more species stabilized their systems. To that end, we analyzed three models of evolution, two predator-prey models and one competition model of evolutionary suicide, to see how the degree of saturation affects the stability of the system. In the predator-prey models, stability is linked to the degree of saturation. Specifically, a fully saturated community will only show stable dynamics, and unstable dynamics occur only when the community is under-saturated. With the competition model, we demonstrate it to be permanently under-saturated, likely showing such extreme dynamics for this reason. Though not a general proof, our analysis of the models provide evidence of the link between community saturation and evolutionary dynamics. Our results offer a possible placement of these evolutionary “pathologies” into a wider framework. In addition, the results concur with previous results showing greater evolutionary response to less biodiversity and clarifies the effect of extrinsic vs. intrinsic non-equilibrial evolutionary dynamics on a community.

## B. Introduction

The development of evolutionary game theory has been critical to understanding the manner by which evolution proceeds. Evolutionary game theory is a mathematical framework which views the traits of a species as its strategies and the environment in which the species lives as the determinant of the rules of the game (Maynard-Smith and Price, 1973; Maynard-Smith, 1982). Evolutionary game theory has shifted scientists’ understanding of evolution from a simple optimization process under which an organism maximizes its fitness against a static, physical environment to a more dynamic optimization in which multiple species co-evolve. Furthermore, it has given scientists a mathematical framework to describe phenotypic evolutionary changes as a complement to population genetics’ description of genotypic changes.

Critical to analyzing evolution as a game is the concept of the evolutionarily stable strategy (ESS). Similar to the Nash equilibrium of classical game theory, it is a solution to an evolutionary game, the equilibrial evolutionary state in which a rare strategy cannot invade and establish within a population. Also like the Nash equilibrium, an ESS is not merely a single strategy but can exist as a mixture of strategies. Under the strategy species concept which defines a species by the strategy it uses, the mixed-strategy ESS can be thought of as a specific assemblage or community of different species. Because of its simplicity, the ESS concept offers a predictable point for the analysis of evolutionary trajectories. Though powerful, it remained an open question as to whether evolutionary dynamics would actually drive the strategies of a species or group of species to an ESS. Is the ESS achievable? When operating with discrete strategies (i.e., matrix games), it was shown that with replicator dynamics the ESS was an attractor (Taylor and Jonker, 1978; Zeeman, 1980; Zeeman, 1981). It seemed that the ESS was the eventual endpoint of any evolutionary dynamic.

Evolutionary game theory was generalized to continuous trait dynamics which led to new problems and solution concepts for the ESS (Vincent and Brown, 1984). It was shown that with continuous trait dynamics, the ESS would not necessarily be an attractor (Takada and Kigami, 1991). Instead, situations could arise in which the fitness maximum for a species was not a convergent stable (Eshel and Motro, 1981; Eshel, 1983; Abrams et al., 1993; Apaloo et al., 2009). This coincided with studies showing unstable evolutionary dynamics such as Red-Queen dynamics and evolutionary suicide (Rosenzweig et al., 1987; Marrow et al., 1992; Matsuda and Abrams, 1994; Dieckmann et al., 1995; Cortez, 2016). The development of other stability concepts such as convergence stability, neighborhood invasion stability (NIS), and mutual invasability were able to explain some of the results and were incorporated into a larger framework along with the ESS (Brown and Pavlovic, 1992; Abrams et al., 1993; Metz et al. 1995; Gertiz et al., 1998; Apaloo et al., 2009). Other results, particularly those of non-equilibrial dynamics, remain unexplained by the new framework.

A potential resolution may be that the communities analyzed are under-saturated. Ripa et al. (2009) in a predator-prey model noted that under-saturated communities showed signs of instability but with increasing species number and an approach to the ESS, the system recovered stability. Many of the studies on non-equilibrial dynamics are done with only a few species, typically 1 or 2, and did not determine whether the species were at or would have gone to an ESS. In several of these examples, it may be that there are not enough species for the community to be at ESS. The community may simply be under-saturated. If it contained close to or the same number of species as the ESS, the ecological (changes in population sizes) and evolutionary dynamics (changes in the frequency of strategies) might lead to a convergent stable evolutionary equilibrium.

To this end, we analyzed three different models of evolutionary dynamics, two models of predator-prey dynamics and one model of competitive dynamics, to see if communities farther from the ESS in terms of species number are more likely to show unstable evolutionary dynamics compared to communities closer to the ESS. By using the full suite of adaptive dynamics, we can see how increases in species number change the evolutionary dynamics of the system. We use these models for illustrative purposes only with no claim of a general proof. We offer a hypothesis and obtain evidence for its feasibility. We hypothesize that non-equilibrial evolutionary dynamics are likely for models in which the number of species is below that of the ESS and that the unstable dynamics shift to stable evolutionary dynamics as the number of species approaches that of the ESS. This result would provide a unified explanation for some of the disparate evolutionary dynamics seen for continuous trait, multi-species games such as convergent stable dynamics to the ESS, limit cycles and other non-equilibrial dynamics, and evolutionary suicide. In what follows, we first examine our hypothesis in two predator-prey models of coevolution (Brown and Vincent 1992, Dieckmann et al. 1995) and then in a competition model of coevolution (Matsuda and Abrams 1994).

## C. Evolutionary Dynamics in Two Predator-Prey Models

To determine the link between community saturation and stability of evolutionary dynamics, we first analyzed two predator-prey models of co-evolution. Predator-prey systems are known to show unstable and non-equilibrial evolutionary dynamics, especially oscillatory dynamics as predators evolve to maximize predation while prey evolve to minimize it (Rosenzweig et al., 1987; Marrow et al., 1992; Dieckmann et al., 1995; Cortez, 2016). The first model analyzed was from Dieckmann, Marrow, and Law (1995), hereafter referred to the DML model. In their paper, the authors analyzed the evolutionary dynamics of a system consisting of 1 prey species and 1 predator species. They found unstable evolutionary dynamics and they determined the conditions that gave rise to them. The second model comes from Brown and Vincent (1992), hereafter referred to as the BV model. Unlike the Dieckmann et al., Brown and Vincent focused on the number of prey and predator species at the ESS and not on the evolutionary dynamics. Through bifurcation analysis, they measured how varying predator specialization affects the number of species in the community and the distribution of their traits. These models were created and analyzed with different goals in mind. Demonstrating that both show the same link between community saturation and unstable evolutionary dynamics would reconcile their approaches and results. We hypothesize that the unstable dynamics of the DML model result from under-saturated communities and adding species will engender stability in the system, and we hypothesize that the BV model will give unstable evolutionary dynamics when the number of co-evolving species is less than the number at the ESS.

We examined the link between stability and community saturation by determining the evolutionary stability of communities created by the two models with varying degrees of saturation/under-saturation. A community is under-saturated if it has fewer species than the ESS (defined in Apaloo et al., 2009). We measure the degree of under-saturation as roughly the difference between the community’s current species richness and the species richness of the model’s ESS (a greater number indicating a greater degree of under-saturation). We used linear stability analysis to determine the asymptotic stability of the evolutionary dynamics (hereafter stability refers to asymptotic stability). For the predator-prey models, we can vary the species richness of the community, and then examine the stability properties of the co-evolutionary dynamics of the species’ strategy values. By evaluating the Jacobian matrix at a community’s co-evolutionary equilibrium, we can characterize the dynamics as non-oscillatory stable (attracting point), oscillatory stable (attracting cycle), oscillatory unstable (repelling cycle), non-oscillatory unstable (repelling point) based on the dominant eigenvalue.

We are interested in how evolutionary dynamics may become unstable in under-saturated communities. Hence, to ensure that any results were not driven by non-equilibrial population dynamics, population sizes were set to their equilibrium abundances based on the current strategy values found among the species of the co-evolving community. This amounts to a fast-slow assumption for the ecological and evolutionary dynamics, an assumption is typically made in models of adaptive dynamics (Geritz et al., 1998).

To define the multispecies evolutionary game, we use the G-function notation of Vincent and Brown (1987, 2005). The population growth rate of species *i* is defined as

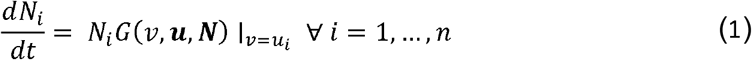

where *G*(*v*, ***u, N***) is the fitness (as defined by per-capita growth rate) of a focal individual with strategy *v*, ***u*** = (*u*_1_,…, *u_n_*) is the vector of strategies found among the *n* species in the community, and ***N*** = (*N*_1_,…, *N_n_*) is the vector of population sizes for each of *n* species. This fitness generating function, *G*(*v*, ***u, N***), generates the fitness function of species *i* when *v* is set equal to *u_i_*.

By assuming that population sizes are always at their equilibrium, we can set *G*(*v*, ***u, N***) = 0 to find the vector of equilibrium population sizes: 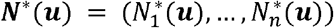. We only consider those species in the community that persist at a positive equilibrium population size. Hence, for all *n* species, 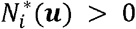.

The strategy dynamics of each species occurs on the adaptive landscape, a plot of *G* versus *v* for the current value of (***u, N***\*(***u***)). We let strategies evolve up the fitness gradient defined by 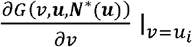. for each *v*. This fitness gradient is evaluated with respect to how an individual’s fitness would change were it to unilaterally change its strategy (the essence of game theory). This defines the fitness gradient as how *G* changes with respect to *v*, the strategy of the focal individual. The evolutionary rate of change of each species’ strategy can be given as (Vincent et al. 1993):

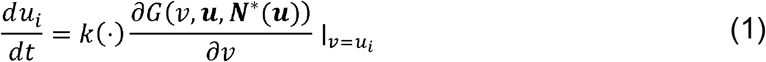

where 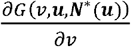 is the slope of the adaptive landscape and *k* represents an evolutionary speed term which may be relatively constant, e.g. a measure of additive genetic variance as assumed in quantitative genetics (Fisher, 1930; Lande, 1982; Falconer and Mackay, 1996), an increasing function of mutation rates, or linear in population size as in the canonical equation of adaptive dynamics (Dieckmann and Law, 1996). An evolutionary equilibrium for the *n* species occurs when 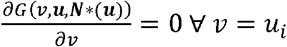, *i* = 1,…, *n*. The corresponding Jacobian evaluated at this evolutionary equilibrium describes the stability of the strategy (evolutionary) dynamics.

### 1. Dieckmann, Marrow, and Law

We begin our analysis with the DML model and write it as a two G-function system, one for the prey species, *G_N_*, and one for the predators, *G_P_*:

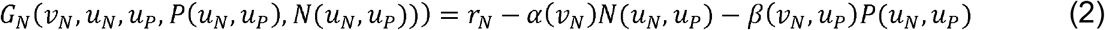

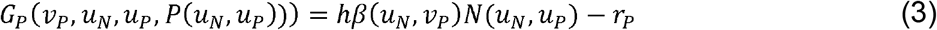

Under our notational scheme, *N* and *P* as subscripts reference the respective term for prey and predator while they refer to the population sizes when written in regular script.

In the DML model, neither the prey nor the predator experience intraspecific frequency-dependent selection, and the predators additionally do not experience intraspecific density-dependence. A predator’s fitness is limited by the population size of the prey and influenced by the traits of the prey and the trait of the focal predator individual. The function *α*(*v_N_*) determines the strength of density dependence of the prey while *β*(*v_N_, v_P_*) determines per-capita predation rate. The parameters *r_N_* and *r_P_* give the intrinsic growth rate and death rate of the prey and predator respectively, while *h* is the conversion efficiency of prey consumed into predators. In the original paper, the authors selected parameters such that *α*(*v_N_*) is symmetrical about *v_N_* = 0.5 and also reaches a minimum at that point; *β*(*v_N_, v_P_*) is broadly a matching strategy where predation rate is maximized when *v_N_* = *v_P_*.

The original authors were interested in how their model generated non-equilibrial evolutionary cycling, and as such, only analyzed a community with one prey and one predator. The evolutionary dynamics were simulated under a polymorphic stochastic model, a monomorphic stochastic model, and a monomorphic deterministic model using the canonical equation of adaptive dynamics to derive the evolutionary speed term *k*(·) (equation 2). Furthermore, the authors performed a bifurcation analysis with the monomorphic deterministic model to determine the stability of strategy dynamics. The parameters used in the bifurcation analysis were the ratio between prey and predator evolutionary speeds 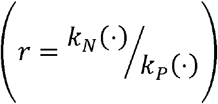 and the conversion efficiency *h*. With this last analysis, they showed that unstable dynamics were more likely to happen at high conversion efficiencies and a high ratio of evolutionary speeds.

For our study, we expand the equations to permit multiple prey and predator species:

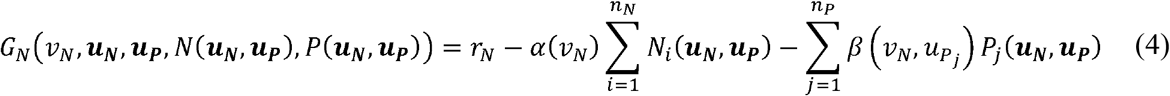

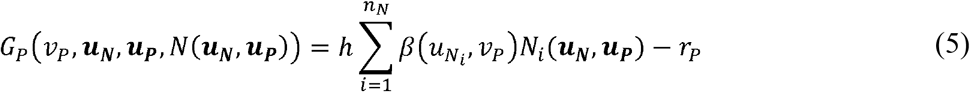

Here, equation (5) denotes fitness of a prey individual with strategy *v_N_* and equation (6) denotes the fitness of a predator individual with strategy *v_P_*. Substituting the strategy of a focal prey individual *v_N_* with a the strategy of species *i*(*u_N_i__*) gives the per-capita growth rate of the *i*-th prey species where *i* = 1,…*n_N_*, and substituting the strategy of a focal predator individual *v_P_* with a the strategy of species *j*(*u_P_j__*) gives the per-capita growth rate of the *j*-th predator species where *j* = 1,…*n_P_*. This generalized form of the model offers not only multispecies dynamics but also allows for speciation at evolutionary branching points (Geritz et al., 1998). Such branching points occur when natural selection (evolutionary dynamics) drive a species strategy to a convergent stable minimum (Brown and Pavlovic, 1992; Abrams et al., 1993) as determined by the second derivative of the per-capita growth rate function. With this multi-species extension, we too performed bifurcation analyses to examine the stability of the evolutionary dynamics in communities that had fewer and/or the same number of prey and predator species as the saturated community. Of particular interest is whether the evolutionary cycling observed in DML resulted from an under-saturated community. Does the non-equilibrium cycling of strategy dynamics disappear with additional prey and predator species? Because we are only interested in the effects of community saturation on evolutionary dynamics, we disregarded the changes in the evolutionary speed term of DML (given by the canonical equation of adaptive dynamics) by setting them constant and equal *k_N_*(·) = *k_P_*(·) = *k*, i.e. *r* = 1. We explored the effect of the conversion efficiency on the number of prey and predator species at the ESS (saturated community), and the nature of the evolutionary dynamics in the saturated and under-saturated communities. In our analysis, we let *h* range along the interval [0, 1] as conversion efficiency was defined as a proportion.

Our analysis of the 1 prey, 1 predator system gives much the same results as Dieckman et al.’s analysis (1995). In order to have a non-zero predator population, the conversion efficiency must be greater than 5 percent, *h* > 0.05. Beyond this point, there is a single trait equilibrium with both prey and predator at *u_N_* = *u_P_* = 0.5. At this point the prey species maximizes its carrying capacity (minimizes the strength of density dependence) and the predator maximizes its fitness by matching the prey’s strategy. As *h* increases, we move from a fully stable attractor to an attractor with asymptotically stable oscillations which occurs just before *h* = 0.08 (Fig. 12a). Moving into greater values of *h*, we see unstable oscillations when *h* ≈ 0.107 (Fig. 12b). Above *h* = 0.13, we get two new equilibrium points symmetrically arranged about *u_N_* = *u_P_* = 0.5. At or slightly above *h* = 0.13, the evolutionary dynamics are visually similar to the Lorenz system as the prey species draws near to one of the new equilibrium points before rapidly evolving to the other (Fig. 12c). At about *h* = 0.181, the evolutionary dynamics become a repeller for the equilibrium *u_N_* = *u_P_* = 0.5 but a locally stable oscillator for the other equilibrium points (Fig. 12d). This holds for *r* = 1.

**Figure 1.**
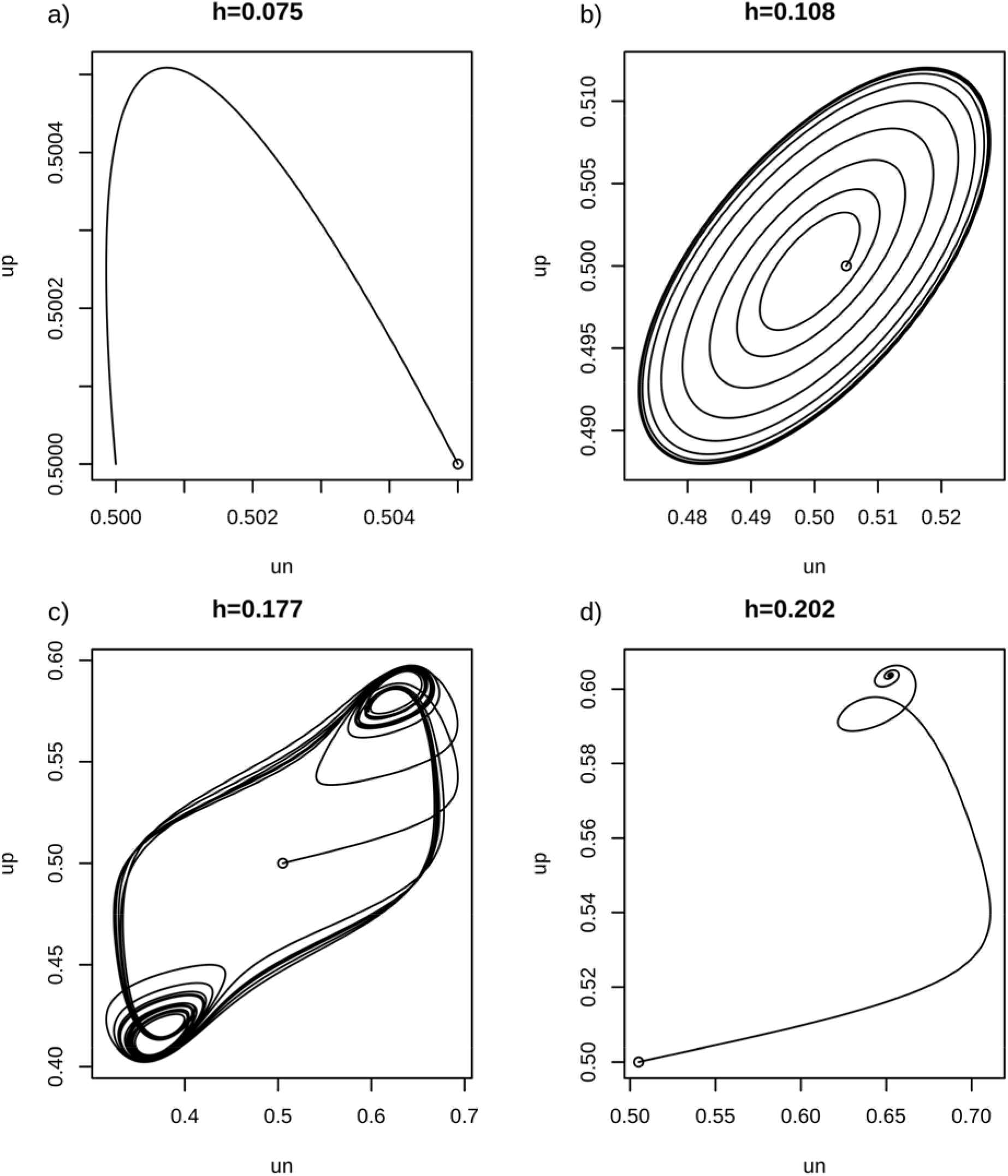
Four types of evolutionary dynamics for a one prey, one predator community over a range of values for *h*. Each graph is a phase portrait of prey evolution (change in strategy) against predator evolution. Open circles represent the initial point of each dynamic with (*u_N_*, *u_P_*) = (0.505, 0.5) a) When *h* = 0.075, the system is stable and returns to the equilibrium. b) When *h* = 0.108, the system is asymptotically unstable with limit cycles. c) When *h* = 0.177, the system is unstable and shows Lorenz-like dynamics around two other equilibria. d) When *h* = 0.202, the system is unstable with regard to the (0.5,0.5) equilibrium and is driven to one of the other equilibria.

Examination of the prey’s adaptive landscape is instructive. When plotting *G_N_* versus *v_N_* for *u_N_* = *u_P_* = 0.5 and *N*\* and *P*\*, we see a switch from a maximum at *v_N_* = 0.5 on the adaptive landscape to a minimum (evolutionary branching point) when *h* = 0.098. Between 0.05 < *h* < 0.098, *u_N_* = *u_P_* = 0.5 is a convergent stable ESS, and one prey and one predator species represents the saturated community. In DML’s original paper, as prey evolutionary speed drastically increases relative to predator evolutionary speed (*r* goes to infinity), the range of *h* under which there are damped oscillations to a stable equilibrium point (with one prey and one predator) asymptotes to the interval (0.098, 0.148). This is noteworthy because the lower bound, *h* = 0.098, occurs at the same value of *h* where the community becomes under-saturated.

In the 1 prey, 1 predator system, the prey is at an evolutionary branching point when *h* > 0.098. We can allow speciation to generate a 2 prey, 1 predator system. The 2 predator, 1 prey system becomes significantly harder to analyze for stability. We can numerically obtain strategy equilibria for prey 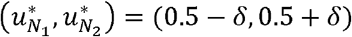 which are symmetric about the predator’s equilibrial strategy values 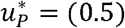. We can also obtain an analytic solution for predator and prey population equilibrium values, but inputting their equilibrial strategy values gives us a divide by 0 error. Without a general solution, we cannot obtain a Jacobian and do linear stability analysis (we can obtain a specific solution to the population equilibrium by first plugging in the equilibrium strategy values, but this cannot be used to determine the Jacobian). Analyzing the adaptive landscapes to determine the stability when the species are off equilibrium is also difficult. When a single species is off their strategy equilibrium (the other two species at strategy equilibrium), its equilibrial population size is negative.

Instead, we analyze stability by a different means. We simulate the population and strategy dynamics of the community by way of the Runge-Kutta 4^th^ order method (Fig. 13). To decouple population dynamics from evolutionary dynamics, we run just the population dynamics for 100 timesteps before running two timesteps of the strategy and population dynamics, allowing the species to get close to the equilibrium population values (we also assume an evolutionary speed term of *k* = 0.1 to slow the dynamics further). We see whether, under our simulations, evolutionary dynamics converge to their equilibrium point or remain oscillatory perpetually. This method is not that extensive (we only simulate for 3 values of *h* = [0.6,0.665, 0.8], 1 value of *k*, and are limited in the specific disturbance from equilibrium) but can give us a patchwork of stability from which we may see patterns. When *h* = 0.6, we see damped oscillations and a convergence back to the equilibrium values. When *h* = 0.8, we see damped oscillations but to a limit cycle (we feel our simulation was run long enough that the limit cycles represent a long-term state of the system). Crucially, when *h* = 0.665, we see damped oscillations to the equilibrium, similar to when *h* = 0.6. This is crucial because the system becomes under-saturated when *h* = 0.665 (a new predator species can invade). It suggests that the flip from a convergent stable system to a convergent unstable system happens when the system is under-saturated. Therefore, it seems that when the community is saturated, it is convergent stable and that only when it is under-saturated can there be convergent instability. While not a true proof, this result tantalizes at a broader picture.

When *h* = 0.665, the predator now exists at a minimum of its adaptive landscape. Being at the minimum means the predator can speciate, creating a 2 prey, 2 predator system. The irregularities of the 2 prey, 1 predator system no longer exist, the Jacobian can be derived, and the 2 prey, 2 predator system can be analyzed through linear stability analysis. We see in this system a simple result, that it is saturated and a convergent stable ESS over the rest of the range of *h*. All species exist on the maxima of their adaptive landscape and all dynamics are stable.

**Figure 2.**
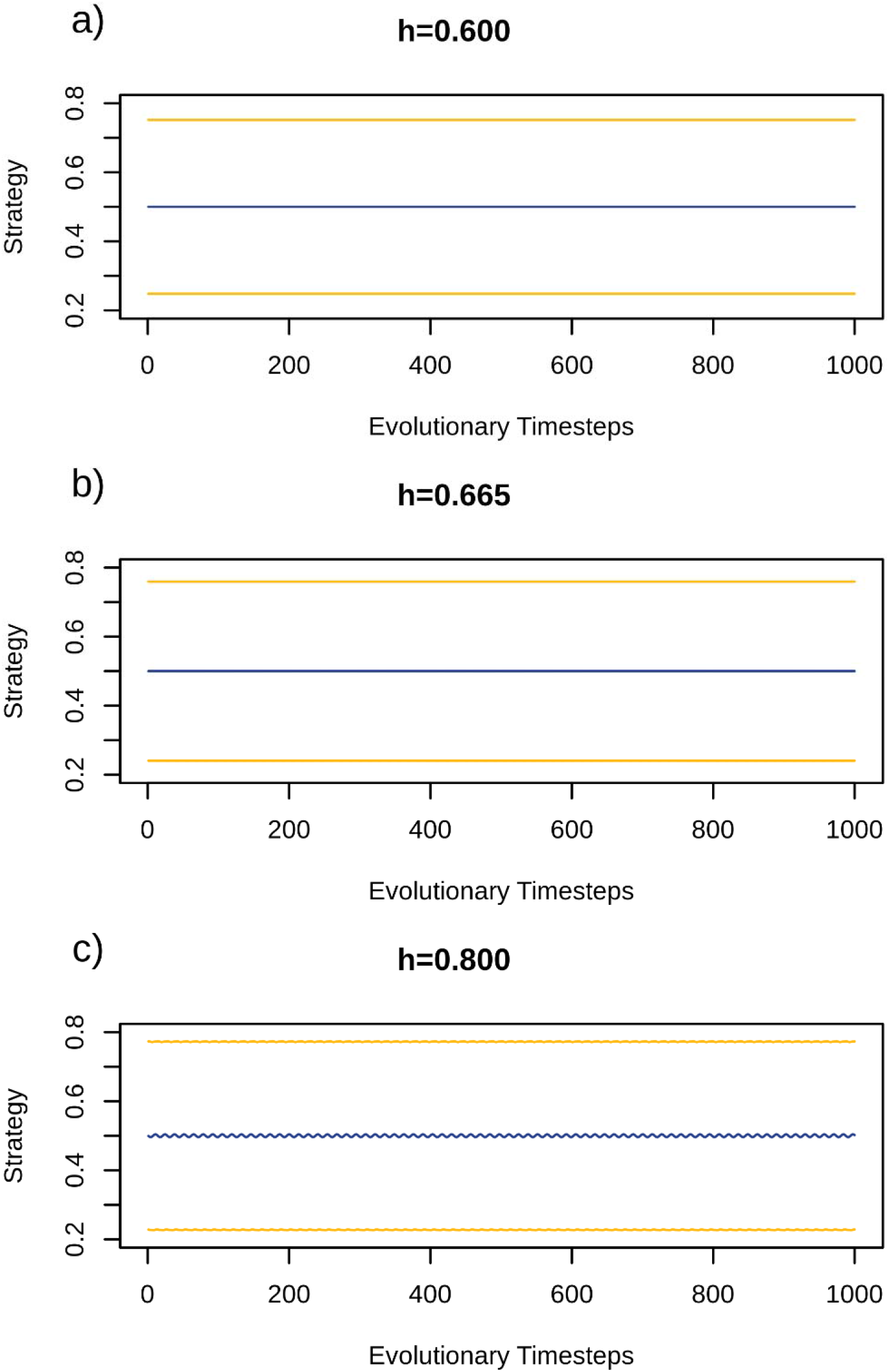
The evolutionary dynamics of a two prey, one predator system under three values of *h*. Gold lines are prey strategy and blue lines are the predator strategy. For each simulation, 100 timesteps of population dynamics were run before the two timesteps of evolutionary dynamics occurred. This was done for 5*x*10^4^ timesteps with 1*x*10^4^ timesteps of evolutionary dynamics. We only show the last 1000 evolutionary timesteps for each run to show the final state of the dynamics. The prey with the smaller strategy value was started at a value 0.02 smaller than its equilibrium and the predator was started at a value 0.01 greater than its equilibrium for all runs. a) When *h* = 0.6, the community is at ESS and the dynamics are stable. b) When *h* = 0.665, the community is not at ESS and the dynamics are stable. c) When *h* = 0.8, the community is not at ESS and the dynamics are unstable showing possible limit cycles.

A very regular pattern of evolutionary dynamics occurred. At low values for the bifurcation parameter *h* (the conversion efficiency of predators), the ESS community has just one prey species and no predator species. The prey’s ESS value of *u_N_* = 0.5 is asymptotically stable. At increasing values of *h*, the ESS becomes a 1 prey, 1 predator community. At first, this community shows asymptotically stable evolutionary dynamics followed by damped oscillations (Fig. 14a). At still higher values of *h*, the one prey, one predator community is no longer an ESS and the prey species is now at a convergent stable minimum (Fig. 14d). At this point, the ESS would be a 2 prey, 1 predator comunity. If the community is forced to remain under-saturated, the convergent stable minimum gives way to unstable oscillatory evolutionary dynamics at ever higher values of *h* (Fig. 14a). With a 2 prey, 1 predator community, stability is recovered. As with the 1 prey, 1 predator community, increasing values of *h* lead to the community no longer being at ESS, this time with the predator at the minimum (Fig. 14d). Once again, though not confirmed through analytic results, it seems that forcing the community to remain undersaturated as *h* increases leads to instability (Fig. 14b, Fig. 13). As *h* goes towards its maximal value of *h* = 1, the ESS is a 2 prey, 2 predator s community. From here on, the community is always saturated and always stable (Fig. 14c and Fig. 14d).

Our analyses of the DML model suggest that the unstable evolutionary dynamics result from under-saturated communities. In all 3 communities – 1 prey, 1 predator; 2 prey, 1 predator; 2 prey, 2 predator – stable dynamics were seen when the community was at ESS while unstable dynamics only appeared when the community was under-saturated (Fig. 14).

**Figure 3.**
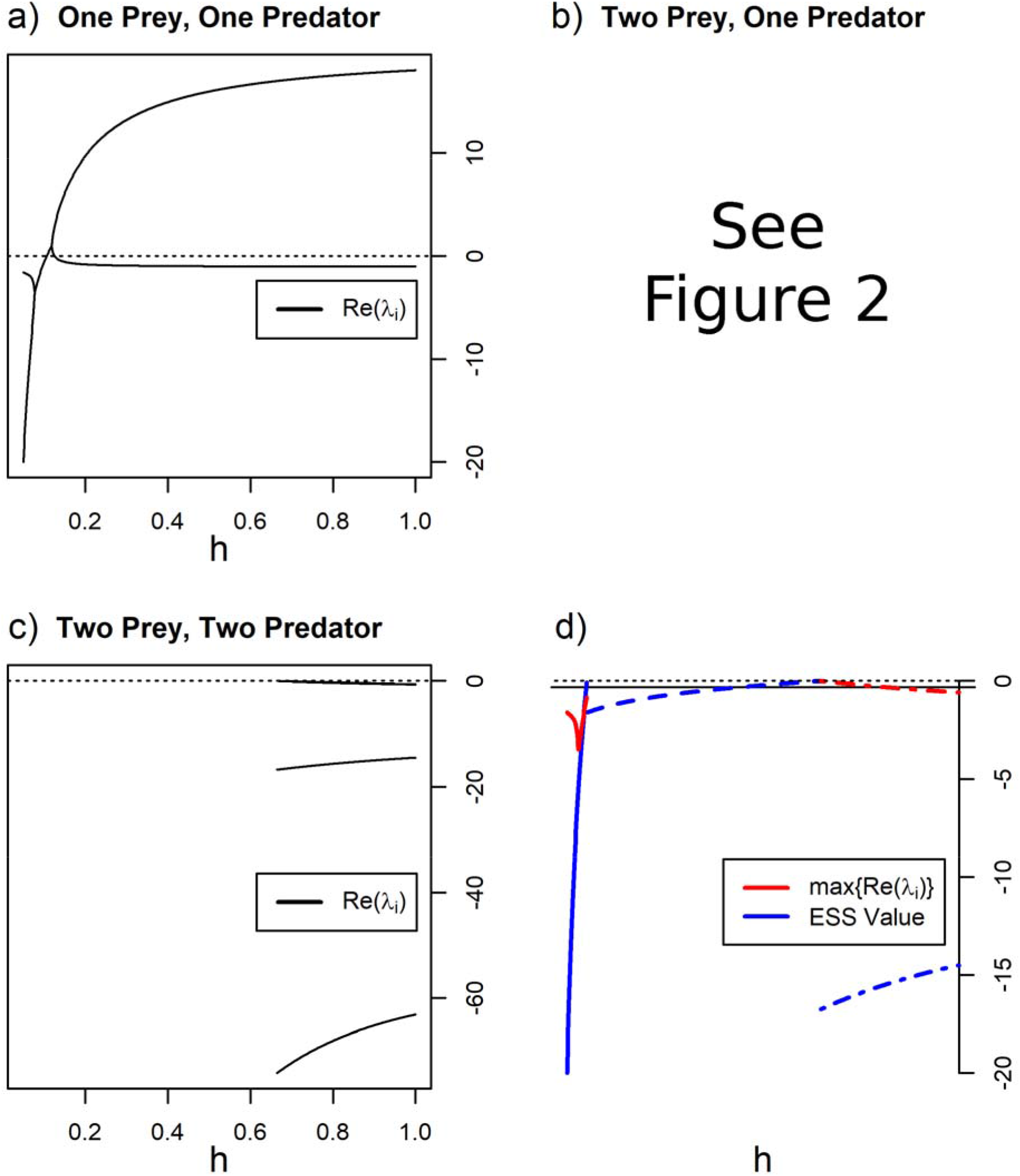
The full analysis of the DML model. a) The real parts of the eigenvalues of a one predator, one prey community over the full range of *h*. The community transitions from non-oscillatory stable to oscillatory stable to oscillatory unstable to non-oscillatory unstable. b) The eigenvalues of the Jacobian for the two prey, one predator system at ecological equilibrium could not be solved for. c) The eigenvalues for the two prey, two predator community. It is always non-oscillatory stable. d) The ESS value (blue) and maximum of the real parts of the eigenvalue (red) for each respective community. Solid lines are the one predator, one prey; dashed lines are the two predator, one prey; and dotted-dashed are the two predator, two prey. We only show when the community is saturated, and in all cases for which the eigenvalues could be solved for show stability with the maximum of the real parts of the eigenvalues being negative.

### 2. Brown and Vincent

We now examine the model from Brown and Vincent (1992). The model was originally a discrete time model, but we convert and analyze it as a continuous time model. As noted in Vincent and Brown (2005), the ESSs of discrete and continuous time forms of G-functions remain the same. Like DML, there is one G-function for the prey species and one for the predator species:

Prey:

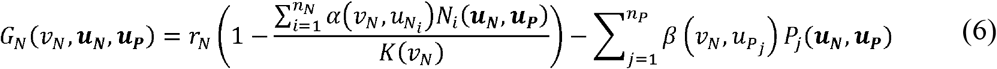

Predator:

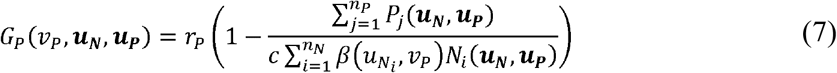

Unlike the DML model, there is intraspecific frequency-dependent selection among prey populations and direct intraspecific density dependence among the predators. Both prey and predator trait space (*v_N_* and *v_P_*) correspond to the domain of real numbers. In this model, the function *α*(*v_N_*, *u_N_i__*) determines competition between prey species based on trait similarity – the more similar the strategies, the greater the competition – *K*(*v_N_*) is a unimodal Gaussian function which determines the carrying capacity of the prey species based upon its strategy *v_N_*, and *β*(*v_N_*, *u_P_j__*) is the capture efficiency of the predator where capture efficiency is maximized when the prey and predator strategies match. The term *c* is the conversion factor for prey consumed into predator offspring.

In the original paper, the authors were interested in how varying the degree of predator specialization *σ_P_*, located in *β*(*v_N_*, *u_P_j__*), influenced species richness and community structure. Therefore, they focused on the saturated communities at their ESS. They evaluated the second derivative of the G-functions (7)–(8) with respect to *v_N_* and *v_P_* to determine whether candidate solutions are at peaks of the adaptive landscape, a necessary criterion for an ESS.

The degree of predator specialization *σ_P_* is a bifurcation parameter. As shown in BV, the ESSs go from a community with one prey and one predator species at low degrees of specialization to successively higher numbers of prey and predator species as the predator becomes more specialized. For a given level of predator specialization, the community can have *n* or *n* + 1 prey species and *n* predator species. Brown and Vincent (1992) did not analyze strategy dynamics towards or away from these ESS communities. This makes the BV model an interesting one for seeing whether evolutionary dynamics are stable in the saturated ESS communities and whether non-equilibrial evolutionary cycling of strategy values occurs when communities are under-saturated. We analyze the BV model the same way we analyzed the DML model to see whether we get similar qualitative results, the results being that evolutionary cycling can occur in under-saturated communities and increasing community saturation leads to more stable evolutionary dynamics.

Deriving the Jacobian for the 1 prey, 1 predator evolutionary systems (population sizes set to their equilibrium values for the given strategy values), we get the eigenvalues 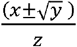 where 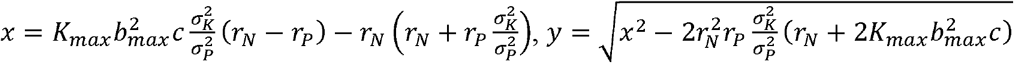 and 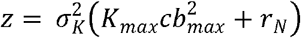. Note that the magnitude of *x* is greater than the magnitude of *y*, |*x*| > |*y*|. Therefore, the sign of the dominant eigenvalue is equal to the sign of *x*. Looking at *x*, it can only be positive – and therefore the system unstable – if *r_N_* > *r_P_* since all parameters are positive.

With the eigenvalues derived, we now did bifurcation analyses similar to the DML model. We varied *σ_P_* from 10 to 0.75, like the original paper, to see how under-saturation affects evolutionary dynamics with a few changes to the original parameters. Firstly, the original parameter set had *r_N_* = *r_P_* = 0.25. Under our stability analysis, this system would always be stable. Therefore, we increased *r_N_* to 0.5. Secondly, even with the increase in *r_N_*, we did not see unstable dynamics over the range of *σ_P_*. Therefore, we increased conversion efficiency from *c* = 0.25 to *c* = 1 to give the possibility of unstable dynamics.

Analysis of the BV system shows the same basic pattern seen in the DML model. Here, the evolutionary dynamics became more unstable as *σ_P_* was lowered. In the 1 prey, 1 predator community, there was only one equilibrium with prey and predator having the same strategy *u_N_* = *u_P_* = 0. With *σ_P_* = 10, this equilibrium was a non-oscillatory attractor. The switch to oscillatory attractor occurred around *σ_P_* = 7.6, to an oscillatory repeller around *σ_P_* = 2.65, and to a non-oscillatory repeller around *σ_P_* = 0.95. The prey also reach an evolutionary branching point at 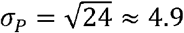. In this case, when the community is at ESS, its dynamics are stable while unstable dynamics only happen when the community is not at ESS and the prey strategy is at a minimum of the adaptive landscape. When at a minimum, there is a range of *σ_P_* for which the dynamics are convergent stable. But, as *σ_P_* becomes smaller and crosses a threshold value, the minimum is not convergent stable and the evolutionary dynamics become unstable.

In the 2 prey, 1 predator system, the equilibrium trait value for the predator remains the same 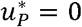 and the prey are now symmetrical around 0, 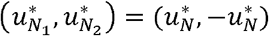. In this community, there are no longer any unstable dynamics over the range of *σ_P_* though there are still oscillatory dynamics with the switch from a non-oscillatory attractor to an oscillator attractor occurring at approximately *σ_P_* = 2.85. One interesting feature is that even after the switch to an oscillator attractor, the dominant eigenvalue pair continues to decline until *σ_P_* = 0.9. One would expect a tendency towards instability with the pair of eigenvalues at some point increasing as was the case for the 1 prey, 1 predator community. This though is only a local phenomenon; beyond *σ_P_* = 0.9, the values of both conjugate eigenvalues increase until they turn positive at *σ_P_* = 0.45 (see Fig. 28, Appendix C). The switch to an under-saturated community occurs before the switch to an unstable dynamics, occurring at approximately *σ_P_* = 2.35. Overall, the pattern was maintained: evolutionary stability when 2 prey and 1 predator species represent the saturated community and non-equilibrial dynamics only when the saturated community has more than 2 prey and 1 predator species.

At 2 prey, 2 predators, we once again see the same pattern. The system is stable over the range of *σ_P_* with the switch to oscillatory dynamics when *σ_P_* = 2.05. Under-saturation occurs around *σ_P_* = 1.4. It is at this point that we stopped our analysis since the 3 predator, 2 prey community could be analyzed neither analytically nor numerically. But, it appears that saturated ESS communities give stable dynamics, and that when prey and/or predator species are at a minimum of their respective adaptive landscapes non-equlibrial evolutionary cycling becomes possible and increasingly likely as the bifurcation parameter is tuned towards increasing species richness at the ESS.

**Figure 4.**
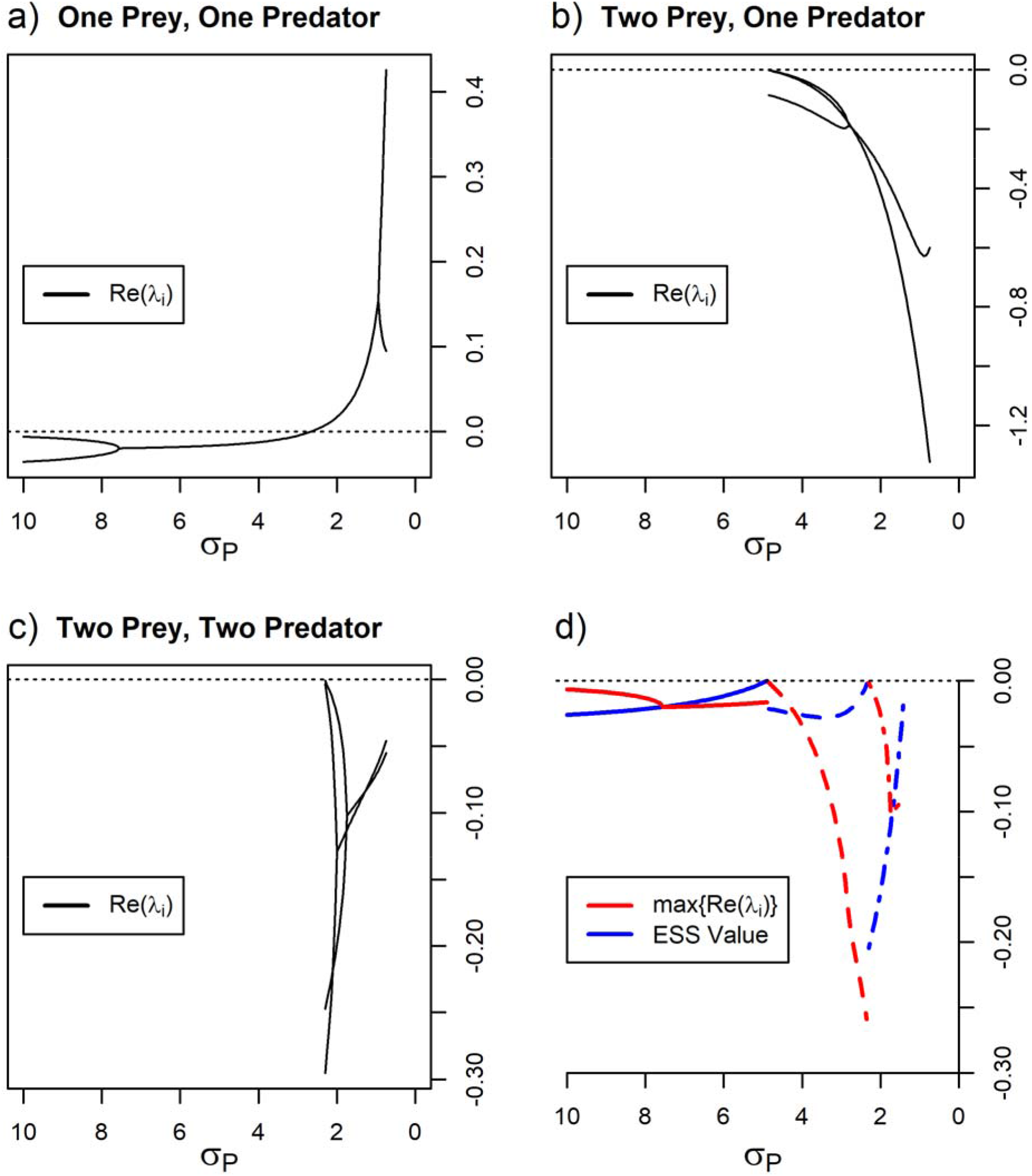
The full analysis of the BV model. The dynamics are nearly identical to the DML model, the only difference being that the two prey and one predator system is analyzed by linear stability analysis (b).

## D. Evolutionary Dynamics in a Competition Model

Our last model derived from Matsuda and Abrams (1994) departs from the previous two. It considers just a single G-function with a community of competing species. We can frame the MA model into the following multi-species G-function:

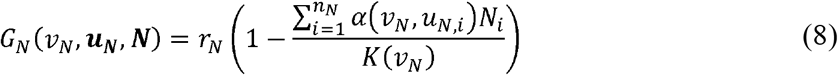

This model is similar to the prey competition portion of the BV model but with two key differences. Firstly, the carrying capacity function is a lognormal function with respect to potential prey strategy values and a lower bound at 0 (only positive strategies allowed). Secondly, there is a term *β* within the competition function which allows for asymmetric competition. If *β* is less than zero, there is a competitive advantage of an individual having a slightly smaller strategy than the rest of the population, and intraspecific frequency dependent selection drives the population to smaller strategies; if *β* is positive, then the opposite occurs.

The original paper showed that when there is only a single species, there is a single globally convergent stable equilibrium with the prey’s strategy value at the maximum of the carrying capacity function if *β* = 0. *β* If is positive but less than a critical value 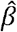, then the original strategy equilibrium increases and a new convergent-unstable equilibrium appears at a value larger than the original equilibrium. This convergent unstable equilibrium divides the strategy space into two domains. If a species has a strategy value less than the convergent-unstable equilibrium, it will move to the original convergent stable equilibrium; if greater, it will increase perpetually to infinity. This leads to evolutionary suicide as the increasing strategy values lead to smaller and smaller population sizes and eventually zero as the strategy value approaches infinity. As *β* increases, the two equilibria move closer together until they are equal; this occurs when 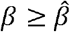. When this happens, the species’ strategy will always evolve towards infinity and evolutionary suicide. Here we ask whether this evolutionary suicide is another manifestation of under-saturated communities.

The original paper did not assess whether a community with a single species was at an ESS, and hence saturated. Graphically representing the adaptive landscape shows that the single species will be at or near an evolutionary branching point regardless of whether it is at the convergent stable equilibrium or evolving to infinity (Fig. 16a). This suggests that the one species community is indeed under-saturated. In fact, simulating the speciation process shows that there can be multispecies communities, each species at a convergent stable equilibrium (Fig. 16b). Examination of the adaptive landscapes under increasing numbers of species suggests that the entire system is permanently under-saturated and could, in fact, be a situation of unlimited niche packing (Roughgarden, 1979; Barabas et al., 2012; Cressman et al., 2017).

To that end, we did a simple exploration of this possibility by constraining the available trait space and seeing how many species would exist at maxima of the adaptive landscape. This trait space would be bounded between 0 and some positive value. To create this finite trait space, we evolutionarily fixed a species at the positive value of trait space. By fixing a species, this acts as a block for other species which cannot evolve beyond that point, effectively acting as a boundary. We gradually increased the trait space (the fixed species’ strategy) and saw how species richness would change as a result. Our results show that as trait space increased, species richness also increased (Fig. 16d). In fact, species richness as a function of available trait space is super linear as seen by the fact that the span under which a specific species number existed decreased (Fig. 16e). We suggest that the MA model is permanently under-saturated and likely an unlimited niche packing model. It may be due to this permanent under-saturation that we see extreme non-equilibrial dynamics that can lead to evolutionary suicide.

**Figure 5.**
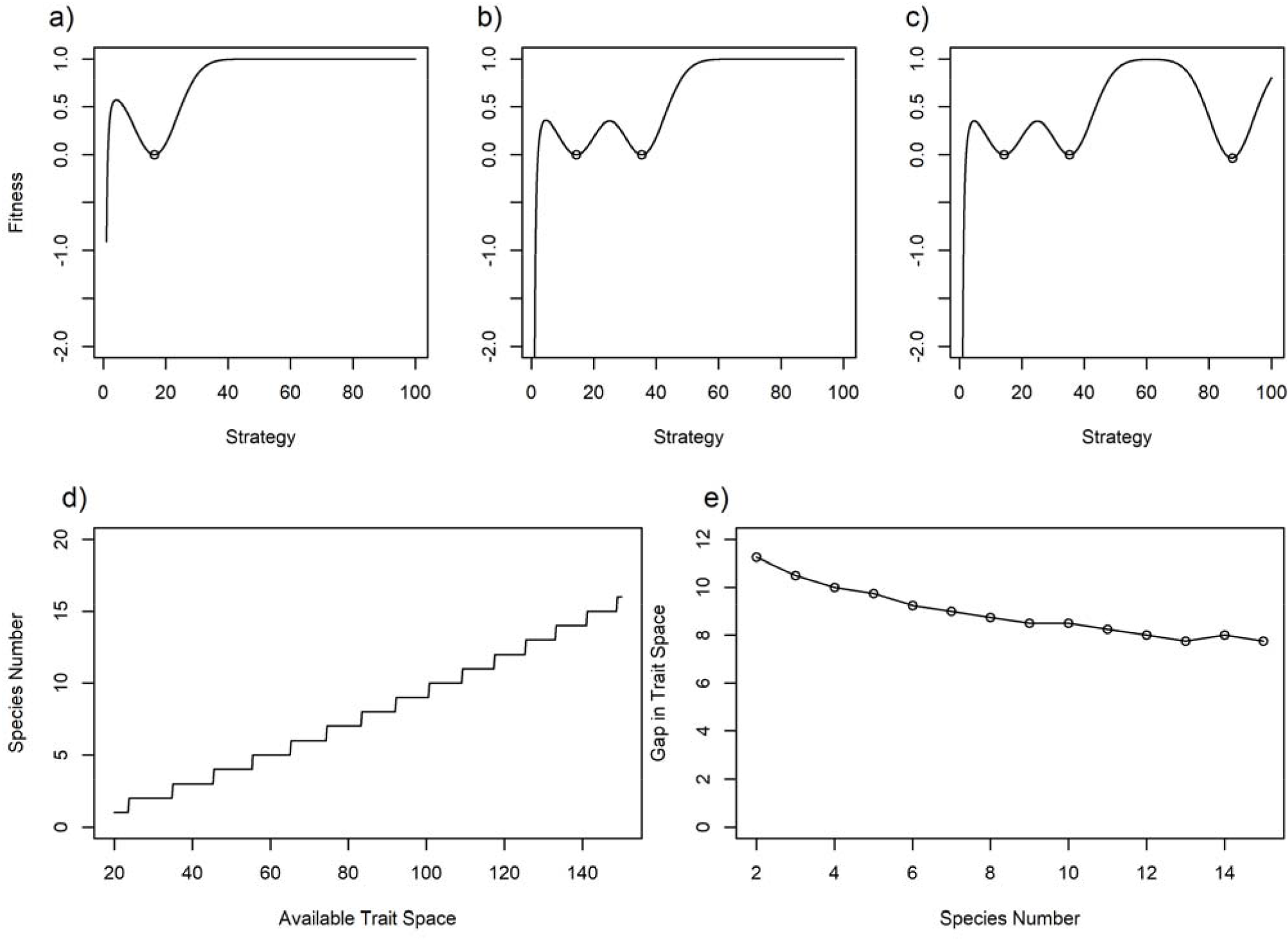
Analysis of the MA evolutionary suicide model. For a-c, we plot the adaptive landscapes with a varying number of species. We select *β* such that there is a domain of convergence and a domain of evolutionary suicide. a) Starting with one species, it converges to a stable minimum. b) A new species arises and both coexist stably at minimums. c) When a third species is added, it rapidly evolves to higher and higher strategy values, i.e. evolutionary suicide. As it evolves to such extremes, the landscape grows behind it allowing for more niche species into which species can invade. The species evolving to evolutionary suicide though is perpetually on the other side of the valley and will never be able to access such niche space. For d-e, we see the number of species that exist at maxima given a specific amount of available trait space. Trait space was constrained by fixing a species evolutionarily at a specific strategy value. d) The number of species that exist at maxima given the available trait space. e) The gap in trait space to the addition of a new species. Each point represents the increase in the amount of trait space needed to get to a certain number of species from the previous number. For example, the first dot indicates that it takes an increase of approximately 11 units of trait space to go from a single species at a maximum to two species at maxima. The decline in the gap between species suggests that species number is super linear with respect to trait space.

## E. Discussion

The ESS concept is an elegant and intuitive solution to an evolutionary game in which no individual can improve fitness by unilaterally changing strategy (Maynard Smith and Price, 1973). By way of the ESS maximum principle, the strategies of the ESS must maximize the fitness of an individual given the circumstances which include the strategies of others (Brown and Vincent 1987). As such, the strategies of the ESS reside at peaks of their adaptive landscape where per capita growth rates are zero if the populations of each strategy have attained equilibrium. Subsequent work revealed other important facets of evolutionary stability. For attainability of the ESS, and even a minimum of the adaptive landscape, there needs to be convergent stability with respect to the eco-evolutionary dynamics (Brown and Pavlovic, 1992; Abrams et al., 1993; Metz et al. 1995). NIS provides an even stronger form of convergence stability for an eco-evolutionary equilibrium point. In this case, the strategy of the ESS or convergent-stable minimum can invade a community that possesses a nearby strategy at its equilibrium population size (Apaloo et al., 2009). Finally, an ESS may exhibit the property of mutual invasibility, a stronger form of NIS where two species with strategies that straddle the ESS can coexist (Geritz et al., 1998). As such a community can possess more coexisting species than an ESS community so long as each species is not at an evolutionary equilibrium. These three additional stability concepts have formed a cohesive framework for understanding equilibria in evolutionary game theory.

Evolutionary games can also exhibit non-equilibrium strategy dynamics. Games with continuous-trait strategies have shown other sorts of such as limit cycles, as noted for variants of the rock-scissors-paper game, and evolutionary suicide (Weissing, 1991; You, 2018). While the four equilibrial properties of the ESS, convergence stability, NIS, and mutual invisibility are well formalized (Apaloo et al., 2009), these non-equilibrium evolutionary dynamics have thus far defied clear context or intuition. They remain evolutionary “pathologies” rather than well-integrated facets of stability properties in evolutionary games.

Ripa et al. (2009) saw unstable evolutionary dynamics in a predator-prey model of species coexistence and coevolution. Using an extension of the BV model which could produce a diversity of competitor species at the ESS even in the absence of predators, they would start with one prey and one predator species to model speciation and community structure. Speciation of either a predator or prey species would occur so long as the species was either not at ESS or showing non-equilibrial dynamics. They noticed that if a community was unstable, it often had many fewer species than the number of species at ESS. By gradually adding species, such communities would transition from an unstable state to a stable state as species number approached that of the ESS. Testing out the Ripa et al., 2009 hypothesis, we examined three evolutionary games using a G-function approach. We examined two predator-prey games and one game of competition. All three models showed unstable evolutionary dynamics only when their communities were under-saturated, and the two models (the predator-prey models) that had finite numbers of species at their ESS showed stable evolutionary dynamics when their communities were saturated. While this is not a proof of the conjecture, the results are very suggestive of a more general phenomenon.

There are two ways for achieving non-equilibrial dynamics in these evolutionary games. First, a community could have non-equilibrial population dynamics. If the optimal strategies are density-dependent, then oscillatory or chaotic population dynamics would produce similar behaviors in the evolutionary dynamics. We removed this possibility by having a timescale separation of fast population dynamics and slow evolutionary dynamics. By setting the species’ population sizes to their equilibrium values given their strategy values, all dynamics were evolutionary. Second, increasing the evolutionary speed term *k*(·) in a multi-species evolutionary game could destabilize the evolutionary equilibria simply by introducing very fast dynamics. While this always remains a possibility for the system, it would be more likely to produce instabilities in the saturated ESS community than in the under-saturated communities due to the interaction between multiple species. Furthermore, the sign of the eigenvalues in our analyses would not change so long as the evolutionary speed term were constant with respect to population dynamics; i.e. were simply additive genetic variance. Due to these modifications, we could be sure that the effects we were seeing were singly a function of community saturation

The results from the predator-prey models gave the most support to our conjecture. With the two models, we did bifurcation analysis on parameters that governed potential diversity while looking at the changes in stability. Both showed that as potential diversity rose while communities remained under-saturated, the communities also became more unstable. In addition, saturated communities were always stable. These results offer a clear link between community saturation and evolutionary stability. This is all the more striking since the two models were structured differently with different goals in mind. The authors of the DML model were interested in evolutionary oscillations which is reflected in the structure of their model with the lack of intraspecific frequency dependence for predators and prey and intraspecific density dependence for predators. The authors of the BV model were interested in community structure and sought equilibrial dynamics reflected in the inclusion of predator intraspecific density dependence. Despite the fact that the models were independently derived and created for separate goals, a similar analysis of them converges on the same result.

One result of note in the BV model occurred in the 2 prey, 1 predator community. In this community, as potentially diversity increased, it became more stable as the dominant eigenvalue became more negative. This even as the system transitioned from non-oscillatory stable to oscillatory stable. We believe it is due to the fact that the system is driven by the predator. One telling fact is that as predator specialization initially increases (*σ_P_* decreases), the equilibrial strategy values for the prey actually move farther apart. The likely explanation is that there is little advantage to prey divergence from maximum carrying capacity if predatory niche breadth is wide. As predatory niche breadth become smaller though, the advantage of divergence of the prey’s strategy values increases and the prey move apart in trait space. This works to the advantage of the predator; prey divergence decreases competition between the prey species and increases individual and cumulative prey population, which then boosts the predator’s population despite decreasing predation rate. Therefore, initial increases in predator specialization benefits the predator. This though is only a local phenomenon because as predator specialization increases, predation rate, and therefore the predator population drops to 0. This is also reflected in the fact that eventually, the dominant eigenvalue rises again and eventually the community becomes unstable.

Evidence from analysis of the MA model also suggests that evolutionary instability results from under-saturated communities. This model shows not only unstable but non-equilibrial evolutionary dynamics under certain conditions. The results of our analysis strongly support the idea that communities from the MA model are permanently under-saturated. As the various species evolve in strategy values towards infinity, the adaptive landscape grows behind them, creating extra niche space available to additional species. Furthermore, species number seems to be a super-linear function of trait space width. The fact that any community will always be permanently under-saturated may be a reason why there is perpetual non-equilibrial evolutionary dynamics. Because of this permanent under-saturation though, we can never see if the community would become evolutionarily stable. Therefore, the evidence from the MA model is less suggestive but still points towards our conjecture.

The MA model seems to have unlimited niche packing (a continuum of an infinite number of species) though evolutionary dynamics for a certain finite number of species can result in the collapse of the system as some of the species evolve higher and higher strategy values resulting in their extinction. It must be noted that this is in contrast to other single function competition models of unlimited niche packing (Gyllenberg and Meszéna, 2005; Meszéna et al., 2005; Szabó and Meszéna, 2006; Parvinen and Meszéna, 2009; Barabas et al., 2012; Barabas et al., 2013; Cressman et al. 2017). For example, in the absence of a predator, the BV model becomes the Roughgarden model (1979). Starting from a finite number of species with distinct strategies, eco-evolutionary dynamics will drive at least some of them to convergent stable minima. Allowing for speciation, the number of species begins to multiply indefinitely. As seen in Cressman et al. (2017), the Roughgarden model does not show non-equilibrial evolutionary dynamics in under-saturated communities. We conjecture the reason for this may have to do with the manipulation of the adaptive landscape. A potential reason that predator-prey models are much more likely to show evolutionary oscillations is that predators are able independently to modify the prey’s adaptive landscape, specifically causing evolutionary minima for the prey. These distortions of the adaptive landscape can cause evolution within the prey as they evolve away from the minimum. It is likely that the extreme asymmetry of competition in the MA model so distorts the adaptive landscape that there is perpetual evolution. Analyzing the dynamics of the adaptive landscape and its interaction with the evolution of a species may be the key to broaden the framework of evolutionary game theory that it can incorporate these evolutionary “pathologies”.

As mentioned before, we have offered evidence for a potential resolution between non-equilibrial evolutionary dynamics and the ESS concept, not a general proof. It remains to be seen whether this is a phenomenon that can be generalized and proven. Analysis of alternative models with distinct features could add to the robustness of our result. For example, with the parameters used for analysis of the BV model, the prey only diversified due to aposematic selection from the predators. Changing the parameters could allow for independent diversification within the prey, likely enhancing community instability (Roughgarden, 1976; Ripa et al., 2009; Cressman et al., 2017). Additionally, asymmetric interactions could be added to the predator capture rate and prey competition within the predator-prey models. Another type of evolutionary instability not analyzed is taxon cycling, the phenomenon in which a species evolves to be later outcompeted by another species with a similar strategy (Rummel and Roughgarden, 1983; Rummel and Roughgarden, 1985; Taper and Case, 1992). Analyzing these and other additional models may bring more evidence for our hypothesis.

More generally, the relationship between biodiversity and non-equilibrial dynamics has received significant attention. Studies have explored the possibility that non-equilibrial dynamics allow for the coexistence of species and the maintenance of hyper-diverse communities (Huisman and Weissing, 1999; Andersen, 2008). Our results suggest the opposite relationship between biodiversity and non-equilibrial dynamics. In our communities, non-equilibrial evolutionary dynamics are a function of community structure, not the other way around, with less diverse communities showing non-equilibrial dynamics. This concurs with previous work that states that biodiversity stalls evolutionary response to environmental change and that rapid evolution is most associated with marginal and newly colonized habitats (Millien, 2006; Mazancourt et al., 2008). This apparent difference is likely due to the nature of the dynamics. Non-equilibrial dynamics driven from external sources like environmental stochasticity are likely to keep the community off an equilibrium and grant species positive population size that would otherwise go extinct. The non-equilibrial dynamics we described though are internally driven, resulting in the need for community under-saturation. It is critical that the distinction between internally driven and externally driven non-equilibrial dynamics is made.

Throughout this paper, we have provided evidence that evolutionary instability is a function of community saturation. That said, we have not formally proved it to be true. Whether this result can be generalized remains an open possibility. We hope this work inspires further investigations of the relationship.

## Cited Literature

Abrams, P.A., H. Matsuda, and Y. Harada. 1993. Evolutionarily unstable fitness maxima and stable fitness minima of continuous traits. Evolutionary Ecology 7:465–487

Andersen, A.N. 2008. Not enough niches: non-equilibrial processes promoting species coexistence in diverse ant communities. Austral Ecology 33:211–220

Apaloo, J., J.S. Brown, and T.L. Vincent. 2009. Evolutionary game theory: ESS, convergence stability, and NIS. Evolutionary Ecology Research 11:489–515

Barabás, G., S. Pigolotti, M. Gyllenberg, U. Dieckmann, and G. Meszéna. 2012. Continuous coexistence or discrete species? A new review of an old question. Evolutionary Ecology Research 14:523–554

Barabás, G., R. D’Andrea, and A.M. Ostling. 2013. Species packing in nonsmooth competition models. Theoretical Ecology 6:1–19

Brown, J.S. and N.B. Pavlovic. 1992. Evolution in heterogeneous environments: effects of migration on habitat specialization. Evolutionary Ecology 6:360–382

Brown, J.S. and T.L. Vincent. 1992. Organization of predator-prey communities as an evolutionary game. Evolution 46:1269–1283

Cortez, M.H. 2016. How the magnitude of prey genetic variation alters predator-prey eco-evolutionary dynamics. The American Naturalist 188:329–341

Cressman, R., A. Halloway, G.G. McNickle, J. Apaloo, J.S. Brown, and T.L. Vincent. 2017. Unlimited niche packing in a Lotka-Volterra competition game. Theoretical Population Biology 116:1–17

de Mazancourt, C., E. Johnson, and T.G. Barraclough. 2008. Biodiversity inhibits species’ evolutionary responses to changing environments. Ecology Letters 11:380–388

Dieckmann, U., P. Marrow, and R. Law. 1995. Evolutionary cycling in predator-prey interactions: population dynamics and the red queen. Journal of Theoretical Biology 176:91–102

Dieckmann, U. and R. Law. 1996. The dynamical theory of coevolution: a derivation from stochastic ecological processes. Journal of Mathematical Biology 34:579–612

Eshel, I. 1983. Evolutionary and continuous stability. Journal of Theoretical Biology 103:99–111

Eshel, I. and U. Motro. 1981. Kin selection and strong evolutionary stability of mutual help. Theoretical Population Biology 19:420–433

Falconer, D.S. and T.F.C. Mackay. 1996. Introduction to Quantitative Genetics, 4^th^ Edition. Prentice Hall, Harlow, England, UK

Fisher, R.A. 1930. The Genetical Theory of Natural Selection. Dover, New York, New York, USA

Geritz, S.A.H., É. Kisdi, G. Meszéna, and J.A.J. Metz. 1998. Evolutionarily singular strategies and the adaptive growth and branching of the evolutionary tree. Evolutionary Ecology 12:35–57

Gyllenberg, M. and G. Meszéna. 2005. On the impossibility of coexistence of infinitely man strategies. Journal of Mathematical Biology 50:133–160

Huisman, J. and F.J. Weissing. 1999. Biodiversity of plankton by species oscillations and chaos. Nature 402:407–410

Lande, R. 1982. A quantitative genetic theory of life history evolution. Ecology 63:607–615

Matsuda, H. and P.A. Abrams. 1994. Runaway evolution to self-extinction under asymmetrical competition. Evolution 48:1764–1772

Marrow, P., R. Law, and C. Cannings. 1992. The coevolution of predator-prey interactions: ESSs and red queen dynamics. Proceedings of the Royal Society of London B 250:133–141

Maynard-Smith, J. 1982. Evolution and the theory of games. Cambridge University Press, Cambridge, England, UK

Maynard-Smith, J. and G.R. Price. 1973. The logic of animal conflict. Nature 246:15–18

Meszéna, G., M. Gyllenberg, L. Pásztor, and J.A.J. Metz. 2006. Competitive exclusion and limiting similarity: A unified theory. Theoretical Population Biology 69:68–87

Metz, J.A.J., S.A.H. Geritz, G. Meszéna, F.J.A. Jacobs, and J.S. van Heerwaarden. 1995. Adaptive dynamics, a geometrical study of the consequences of nearly faithful reproduction. Pages 183–231 in S.J. van Strien and S.M. Verduyn Lunel, editors. Stochastic and spatial structures of dynamical systems. Royal Netherlands Academy of Arts and Sciences, Amsterdam, the Netherlands

Millien, V. 2006. Morphological evolution is accelerated among island mammals. PLoS Biology 4:e384

Parvinen, K. and G. Meszéna. 2009. Disturbance-generated niche-segregation in a structured metapopulation model. Evolutionary Ecology Research 11:651–666

Ripa, J., L. Storling, P. Lundberg, and J.S. Brown. 2009. Niche co-evolution in consumer-resource communities. Evolutionary Ecology Research 11:305–323

Rosenzweig, M.L., J.S. Brown, and T.L. Vincent. 1987. Red Queens and ESS: the coevolution of evolutionary rates. Evolutionary Ecology 1:59–94

Roughgarden, J. 1976. Resource partitioning among competing species – a coevolutionary approach. Theoretical Population Biology 9:388–424

Roughgarden, J. 1979. Theory of Population Genetics and Evolutionary Ecology: An Introduction. Prentice Hall, Upper Saddle River, New Jersey, USA

Rummel, J.D. and J. Roughgarden. 1983. Some differences between invasion-structured and coevolution-structured competitive communities: a preliminary theoretical analysis. Oikos 41:477–486

Rummel, J.D. and J. Roughgarden. 1985. A theory of faunal buildup for competition communities. Evolution 39:1009–1033

Szabó, P. and G. Meszéna. 2006. Limiting similarity revisited. Oikos 112:612–619

Takada, T. and J. Kigami. 1991. The dynamical attainability of ESS in evolutionary games. Journal of Mathematical Biology 29:513–529

Taper, M.L. and T.J. Case. 1992. Models of character displacement and the theoretical robustness of taxon cycles. Evolution 46:317–333

Taylor, P.D. and L.B. Jonker. 1978. Evolutionary stable strategies and game dynamics. Mathematical Biosciences 40:145–156

Vincent, T.L. and J.S. Brown. 1984. Stability in an evolutionary game. Theoretical Population Biology 26:408–427

Vincent, T.L. and J.S. Brown. 2005. Evolutionary Game Theory, Natural Selection, and Darwinian Dynamics. Cambridge University Press, Cambridge, UK

Vincent, T.L., Y. Cohen, and J.S. Brown. 1993. Evolution via strategy dynamics. Theoretical Population Biology 44:149–176

Weissing, F.J. 1991. Evolutionary stability and dynamic stability in a class of evolutionary normal form games. Pages 29–97 *in* R. Selten, editor. Game Equilibrium Models I. Springer-Verlag Berlin Heidelberg New York Tokyo

You, L. 2018. Spatial and nonspatial evolutionary games and their applications. Thesis, Maastricht University, Maastricht, Netherlands

Zeeman, E.C. 1980. Population dynamics from game theory. Pages 471–497 *in* Z. Nitecki and C. Robinson, editors. Global Theory of Dynamical Systems: Proceedings of an International Conference Held at Northwestern University, Evanston, Illinois, June 18-22, 1979. Springer-Verlag Berlin Heidelberg New York

Zeeman, E.C. 1981. Dynamics of the evolution of animal conflicts. Journal of Theoretical Biology 89:249–270

